# Sialoglycans Modulate Siglec-5 – TLR4 Interactions in Osteoarthritis

**DOI:** 10.1101/2025.03.18.643878

**Authors:** Loise Råberg, Fan Jia, Ula von Mentzer, Vignesh Venkatakrishnan, Niclas G. Karlsson, Alexandra Stubelius

**Affiliations:** Division of Chemical Biology, Department of Life Sciences, Chalmers University of Technology, Gothenburg, SE-412 96, Sweden; Department of Life Sciences and Health, Faculty of Health Sciences, Oslo Metropolitan University Oslo Metropolitan University, Oslo, Norway

**Author notes:** Corresponding author: Alexandra Stubelius, Chalmers University of Technology, Department of Life Sciences, Division of Chemical Biology, Kemigården 4, SE-412 96 Göteborg, Sweden.

**Keywords:** Osteoarthritis, Monocytes, Siglec-5, Sialic Acid, TLR4

## Abstract

Osteoarthritis (OA) is characterized by chronic, low-grade inflammation that contributes to cartilage degradation and joint pain. We previously identified Siglec-5/14 in synovial fluid of OA patients (OA SF), which prompted us to investigate its interaction with the sialylated proinflammatory receptor TLR4 in monocytes. Here, we reveal an inverse correlation between Siglec-5 and TLR4, suggesting Siglec-5 may suppress inflammation. To gain mechanistic insights, monocytes stimulated with OA SFs were compared to M-CSF, LPS, and sialidase, to assess patient-specific inflammatory pathways and phenotypes. Notably, OA SF that triggered elevated IL-6 production in monocytes exhibited phenotypes similar to those of LPS- or sialidase-treated cells, reinforcing the role of sialylation patterns influencing OA severity. To confirm direct interaction of Siglec-5 and TLR4, colocalization was analyzed, displaying a time- and sialoglycan-dependent interaction. These findings reveal a Siglec-5-TLR4 axis modulated by sialylation, highlighting a potential strategy for mitigating inflammation and preserve joint integrity in OA.

## INTRODUCTION

Osteoarthritis (OA) is the most common degenerative joint disorder worldwide, characterized by synovitis, cartilage degradation and subchondral bone remodeling that severely impacts quality of life. ^1^ This complex condition involving mechanical stress and inflammatory pathways displays significant variability between patients. Persistent inflammation stem from both tissue-resident and infiltrating immune cells, including monocytes, which play key roles in perpetuating inflammation. ^2,3^ Upon recruitment, monocytes are polarized into a pro-inflammatory phenotype in response to the local environment, contributing to the cycle of tissue damage and degeneration of cartilage. ^4,5^ The multifactorial nature of OA coupled to the limited regenerative capacity of cartilage has posed a major challenge both for finding relevant biomarkers and developing effective treatments. Despite extensive efforts, clinical trials targeting cytokines such as IL1β and TNFɑ have not yielded significant therapeutic benefits, underscoring the need for alternative mechanistic insights and novel intervention strategies. ^6–8^

Recent advances in nanotherapeutics have offered promising new approaches, as they can deliver drugs to the deeply embedded cells in the cartilage responsible for OA progression. ^9^ In our previous work, we explored the interactions between a family of small dendritic polyamidoamine (PAMAM)-based nanoparticles (NPs) and synovial fluid (SF) proteins from OA patients. ^10^ By proteomics, we identified a high abundance of Sialic acid-binding immunoglobulin-like lectin-5/14 (Siglec-5/14) in the proteins associated with the NPs, suggesting a potential involvement in OA pathogenesis. Despite their opposing immunological roles, Siglec-5 and Siglec-14 share a high degree of extracellular sequence homology, which makes them difficult to distinguish experimentally. ^11^ Siglecs are gaining increasing attention in multiple diseases, including cancer, sepsis, and autoimmune disorders due to their immunomodulatory functions and potential for therapeutic targeting. ^12–14^ Siglec-5 has been found in sepsis, Siglec-9 in arthritis, and Siglec-15 plays an important role in macrophage fusion into osteoclasts but so far, the role of Siglec-5 and Siglec-14 has not been reported in OA. ^15–17^ Through their affinity to sialic acid (Sia), inactivating Siglecs such as Siglec-5 act as key regulators of monocyte activation, recruiting the tyrosine SHP-1 and SHP-2, to counterbalance pro-inflammatory pathways, ^18^ whereas activating Siglecs, like Siglec-14 signal through DAP12. ^11^ Siglec-5 is mostly found on cell surfaces on monocytes, macrophages, and T cells. However, soluble Siglec-5 has been identified and was shown to inhibit CD8^+^ T cell proliferation in sepsis, and soluble Siglec-9 demonstrated immunosuppressive effects on macrophages in arthritis models. ^15,17,19^ The generation of soluble Siglec-14 has been well characterized, where one study demonstrated that it can inhibit the interaction between membrane-bound Siglec-14 and TLR2. ^20^

In arthritis, upregulated Siglec-9 has been identified and murine Siglec-E, the mouse ortholog of human Siglec-9 has been shown to regulate pattern recognition receptors (PRRs). ^15,21^ PRRs, such as toll-like receptor 4 (TLR4), play a key role in OA, its activation can be induced by structural fragments of damaged cartilage. ^22,23^ TLR4 recognizes solubilized fragments of glycoproteins such as collagen type-II, hyaluronan, aggrecan, and lubricin which are present in OA SF and function as damage-associated molecular patterns (DAMPs), thereby exacerbating joint inflammation. ^24–26^ TLR4 is gaining increasing interest as its activation disrupts immune homeostasis and contributes to the onset and progression of age-related diseases, including OA. ^27^ Through the NF-κB pathway pro-inflammatory cytokines such as IL-1, IL-6, IL-8, and tumor necrosis factor (TNFɑ) are released in the joint, aggravating joint degradation and OA symptoms, however its regulation is poorly understood in OA. ^3,23,28–31^ TLR4 activity is influenced by its internalization dynamics and glycosylation decorations.

Specifically, its sialylation patterns play a pivotal role in modulating TLR4 signaling. ^21,32,33^ In general, endogenous activity of sialyltransferases and sialidases are regulated upon monocyte activation and differentiation, and alteration in sialyltransferase expression have been identified in OA cartilage. ^34–36^ It has been shown that LPS can trigger sialidase relocation to the cell surface, where it desialylates TLR4 and promotes its activation. ^37^ Interestingly, Siglec-E counteracts this process by binding Sia residues on TLR4, leading to either internalization or shedding of TLR4 to reduce inflammation. ^21,38–40^ Both Siglec-9 and Siglec-5 have shown high affinity to TLR4, indicating their immunomodulatory role of TLR4 signaling.^32^ Siglec-5 has however not been studied as extensively as Siglec-9, which could be due to lack of a mouse ortholog. ^17,18^ As we had identified Siglec-5/14 in the SF of OA patients, the purpose of this study was to investigate the regulatory relationship between Siglec-5, Siglec-9, Siglec-14, Siglec-15 and TLR4 in monocyte stimulated with OA SF.

We found an inverse correlation between membrane-associated Siglec-5/14 and TLR4 expression by flow cytometric profiling, suggesting that engaging Siglec-5/14 dampens excessive inflammatory responses. To gain mechanistic understanding we employed stimulations including macrophage-colony stimulating factor (M-CSF), IL1β/TNFɑ, LPS, and sialidase to understand patient-specific inflammatory pathways and phenotypes. Further insights of crosstalk between Siglec-5 with TLR4 and Siglec-9, was gained through inhibition of TLR4 signaling using the small molecule inhibitor TAK-242. Our findings suggest that Siglec-5 can serve as a modulator of TLR4-driven low-grade inflammation. High-grade inflammatory conditions induced by LPS or sialidase displayed similar cytokine profiles both with and without TAK-242, indicating that sialoglycans are critical regulators of TLR4-driven inflammation. Overall, our findings suggest that Siglec-5, but not Siglec-9, Siglec-14 and Siglec-15, through Sia expression on cells could present an endogenous regulatory pathway to fine tune and mitigate inflammation to reduce cartilage degradation in OA.

## RESULTS

### Siglec-5/14 can be found in synovial fluid from osteoarthritis patients and membrane-associated Siglec-5/14 negatively correlates with TLR4 expression on monocytes

NPs can be used as enrichment tools to identify proteins associated with pathogenic regulation. In a previous study, we developed a set of NPs which were incubated with OA SF where the proteins were identified through proteomics.^10^ Amongst the proteins found in association with the NPs we identified Siglec-5/14 as one of the 14 most abundant entities (Figure 1A). As Siglec-5 functions as an inhibitory receptor, we used the IntAct database to find a potential regulated partner and discovered a medium relationship (MI score 0.54) between TLR4 and Siglec-5, but not Siglec-14 (Figure 1B), suggesting an unexplored regulatory pathway in OA.^41^ Interestingly, both of these receptors can be found on monocytes, which is the most abundant immune cells in the OA joint .^42^

**Figure 1.**
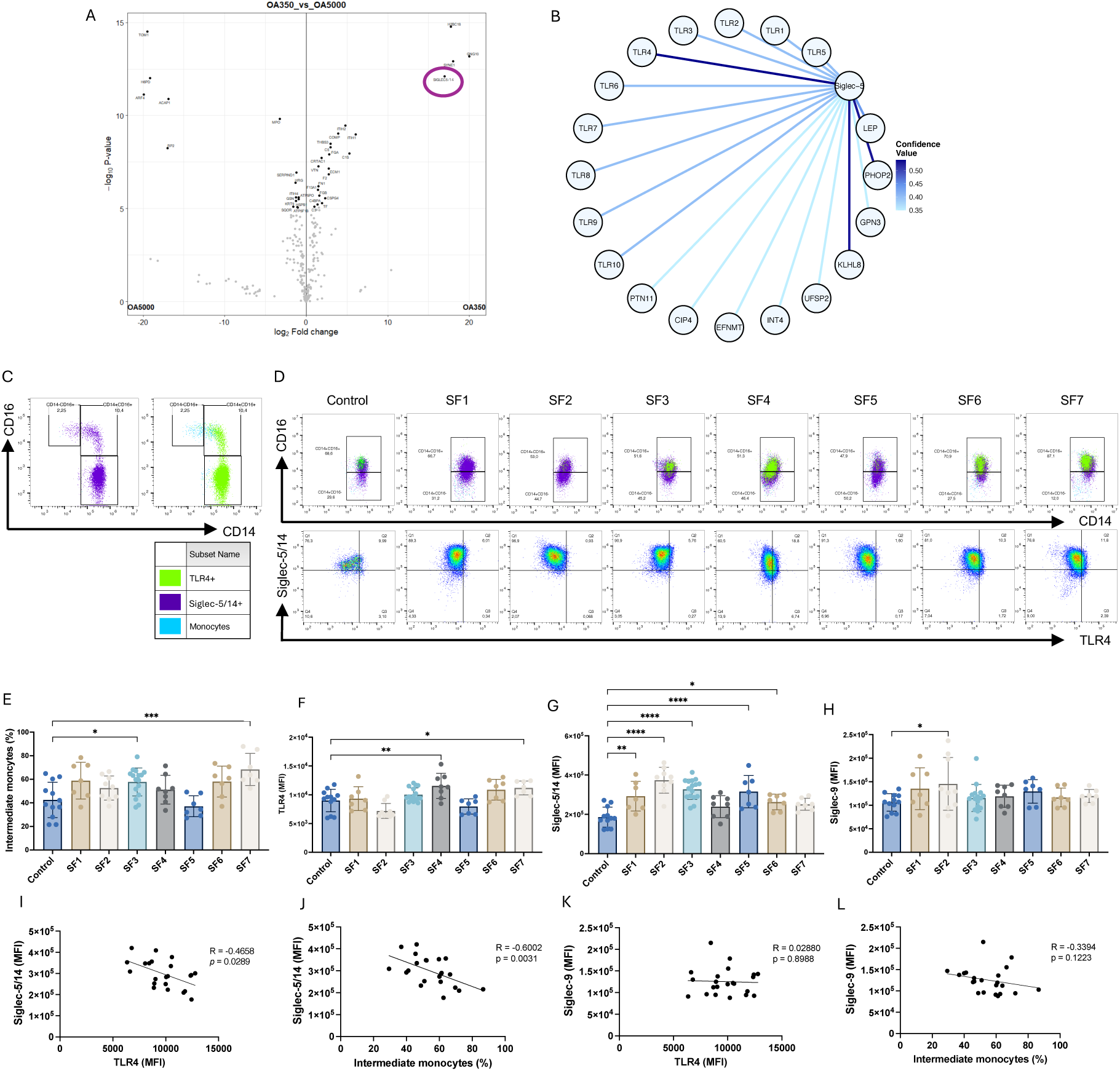
Siglec-5/14 can be found in synovial fluid from osteoarthritis patients and membrane-associated Siglec-5/14 negatively correlates with TLR4 expression on monocytes. (A) Volcano plot displaying the proteins detected in the protein in association with PAMAM_5000_ and PAMAM_350_, here denoted OA5000 and OA350 using proteomics analysis. Siglec-5/14 is indicated with a circle. (B) Protein-protein interaction network visualized from the IntAct database. Medium evidence of interaction appears to TLR4, PHOP2, and KLHL8, where TLR4 had the highest confidence value with a MI score of 0.54. The illustration was adopted from the website and resolution and color scheme improved using R. (C) Representative flow cytometric dot plots displaying the baseline levels (0hrs) of classical, intermediate, and non-classical subsets, where blue color represents all monocytes, purple color represents Siglec-5/14^+^, and green represents TLR4^+^ cells. (D) Representative flow cytometric dot plots of monocytes after 24hrs of coculture with OA SF. The plots display the ratio between classical and intermediate monocytes, where blue represents both subsets, purple represents Siglec-5/14^+^ cells, and green represents TLR4^+^ cells. (E) Percentage intermediate monocytes (CD14^+^CD16^+^) after 24hrs of coculture with OA SF. Each datapoint represents one technical replicate from three independent biological replicates. (F-H) Median fluorescence intensity (MFI) of TLR4, Siglec-5/14, and Siglec-9 on the cell membrane of monocytes after 24hrs of coculture with OA SF. Each datapoint represents one technical replicate from three independent biological replicates. (I-L) Correlation between Siglec-5/14 to TLR4 and intermediate monocytes, and between Siglec-9 to TLR4 and intermediate monocytes. Each datapoint is the mean from three technical replicates from three donors exposed to SF1-7, except SF3 which was evaluated on four different donors. Data is represented as mean±SD, statistical analysis was performed using one-way ANOVA with Dunnett’s multiple comparisons test. Correlation analysis was performed using Spearman’s correlation test (\**p*<0.05; \*\**p*<0.01; \*\*\**p*<0.001; \*\*\*\**p*<0.0001).

Monocyte function can be classified into different subsets based on their expression of CD14 and CD16.^43,44^ The intermediate CD14^+^CD16^+^ subset presents antigens and secretes cytokines and has been found in greater abundance in the joint of OA patients.^42^ In this study, we isolated blood-derived monocytes and confirmed their relative percentage of subsets (Figure S1, Figure S2A and S2B). Figure 1C shows the gating strategy at baseline, where all monocytes were positive for Siglec-5^+^ (purple) and Siglec-9 (not shown), whereas the percentage TLR4^+^ monocytes were mainly found in the classical and intermediate population (green). We found that monocytes cocultured with OA SFs for 24hrs generated an intermediate monocyte population consisting of more than 50% of the population (Figure 1D and 1E). Compared to baseline, the classical monocytes decreased, and non-classical monocytes were completely absent. While most cells remained positive for Siglec-5 (purple), TLR4 (green) expression was reduced.

The monocyte subsets were further analyzed for their baseline expression of TLR4, Siglec-5/14, Siglec-9, and Siglec-15. Classical monocytes displayed the highest expression of TLR4, Siglec-5/14, and Siglec-9, whereas Siglec-15 was not detected on any subset (Figure S2C-E and Figure S3). After 24h of OA SF stimulations, multiple OA SFs led to a significant upregulation in Siglec-5 expression on monocytes (Figure 1G). In contrast to previous murine studies, Siglec-9 was only upregulated in one patient (Figure 1H). Correlation analysis confirmed an inverse relationship between Siglec-5/14 with TLR4 and intermediate monocytes (Figure 1I and 1J), whereas no correlation was observed between Siglec-9 with TLR4 and intermediate monocytes (Figure 1K and 1L). A correlation between Siglec-5/14 to Siglec-9, and intermediate monocytes to TLR4 could not be established (Figure S4A and S4B).

In summary, the monocytes shifted toward the intermediate subset after SF stimulation. Correlation analysis revealed an inverse relationship between Siglec-5/14, TLR4, and intermediate monocytes, but not with Siglec-9, suggesting interactions specifically between TLR4 and Siglec-5 in OA.

### Siglec-5 regulate TLR4 signaling in inflammation

To investigate how specific signaling pathways dictate membrane expression of TLR4, Siglec-5/14, and Siglec-9, blood monocytes were isolated and subsequently stimulated with either M-CSF, IL1β/TNFɑ, or LPS. After 24hrs of stimulation with M-CSF and LPS, an expansion of intermediate monocytes where noted compared to baseline levels. The total monocyte population remained Siglec-5/14^+^, while TLR4^+^ monocytes decreased in the control and LPS (Figure 2A). We further assessed the expression dynamics of TLR4, Siglec-5/14, and Siglec-9 levels at 0, 4 and 24hrs of stimulation (Figure 2B-D). An initial increase in TLR4 could be observed already after 4hrs in the control and M-CSF, which remained after 24hrs.

**Figure 2.**
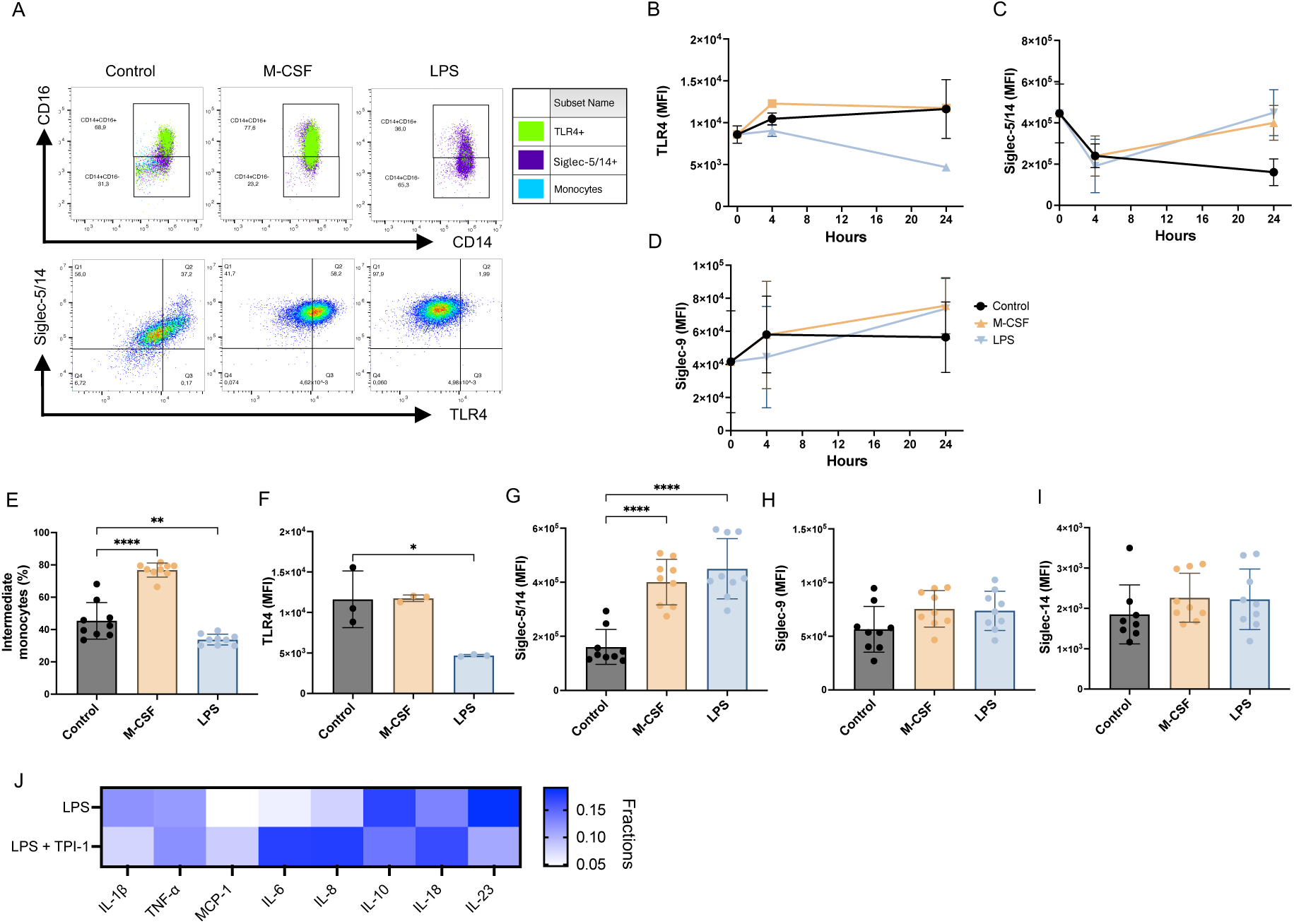
Siglec-5 regulate TLR4 signaling in inflammation. (A) Representative flow cytometric dot plots displaying the percentage classical and intermediate monocytes in the control, and M-CSF- and LPS-stimulated monocytes. Blue represents all cells, green represents TLR4^+^ cells, and purple represents Siglec-5/14^+^ cells, which can also be displayed in the row of dot plots below. (B-D) MFI of membrane-associated TLR4, Siglec-5/14, and Siglec-9 was assessed at 0, 4, and 24hrs of stimulation with M-CSF and LPS and compared to the control. (E) The percentage intermediate monocytes after 24hrs in the control, and monocytes stimulated with M-CSF and LPS. (F-I) MFI of membrane-associated TLR4, Siglec-5/14, Siglec-9, and Siglec-14 in the control and monocytes stimulated with M-CSF and LPS for 24hrs. (J) Cytokines produced by monocytes stimulated with LPS with and without the SHP-1 inhibitor TPI-1 (100ng/mL) was analyzed using the LEGENDplex Human Inflammation Panel 1 (13-plex). Cytokines with levels below the detection limit were excluded from analysis. Each datapoint represents one technical replicate, with three technical replicates performed for each of three independent biological replicates, except for TLR4 which was measured on one biological replicate. Data is represented as mean±SD, statistical analysis was performed using one-way ANOVA with Dunnett’s multiple comparisons test (\**p*<0.05; \*\**p*<0.01; \*\*\**p*<0.001; \*\*\*\**p*<0.0001).

LPS reduced TLR4 expression already within 4 hours and this effect remained after 24hrs. Siglec-5/14 decreased across all groups after 4hrs of stimulation but recovered to basal level expression by M-CSF and LPS stimulations after 24hrs. All stimulations induced Siglec-9 expression after 4 and 24hrs compared to basal levels.

We proceeded with comparing the dynamics of the expression of intermediate monocytes, TLR4, Siglec-5/14, and Siglec-9 at 4 and 24hrs. For most stimulations, intermediate monocytes expanded continuously over 24hrs (Figure S5A, Figure 2E). TLR4 expression increased initially by M-CSF, while LPS reduced the expression of TLR4 compared to control at 4hrs (Figure S5B). After 24hrs, LPS continued to reduce TLR4 expression, whereas M-CSF and IL1β/TNFɑ produced comparable levels to control (Figure 2F, Figure S6A).

Compared to the control, Siglec-5/14 and Siglec-9 expression remained the same across all stimulations at 4hrs (Figure S5C and S5D). After 24hrs of stimulation, M-CSF, IL1β/TNFɑ, and LPS had all induced a significant increase in Siglec-5/14 (Figure 2G, Figure S6B). In contrast, no significant alterations in Siglec-9 expression were observed by any stimulations at 24hrs (Figure 2H). To assess whether the shifts were due to Siglec-5 or Siglec-14, a Siglec-14 specific antibody was used. We found no significant change in membrane expression of Siglec-14 between the control, M-CSF, and LPS (Figure 2I). Siglec-5 mediates its inhibitory immune regulation by recruiting SHP-1 and SHP-2. To confirm that the mechanism involved Siglec-5 rather than Siglec-14, the SHP-1 inhibitor TPI-1 (100ng/mL) was added one hour prior to adding M-CSF and LPS. After 4hrs, supernatants were collected and analyzed for cytokine production using the LEGENDplex assay. LPS-stimulated monocytes produced higher levels of IL1β, IL-10, and IL-23 compared to those treated with TPI-1. In contrast, inhibition of SHP-1 resulted in increased production of MCP-1, IL-6, IL-8, and IL-18 by LPS-stimulated monocytes.

In summary, the intermediate monocyte population expanded over time, where TLR4 was reduced as Siglec-5 increased, displaying an interesting dynamic regulation pattern. This was true for both M-CSF and LPS, whereas IL1β/TNFɑ showed little regulation. Our data suggests complex interacting mechanisms governing TLR4 signaling, ultimately shaping the monocyte response in OA.

### Siglec-5 and TLR4 co-localization on the cell surface of monocytes

Both Siglec-5 and TLR4 are transmembrane receptors which engages extracellularly at the cell-membrane surface as well as interacts with intracellular ligands to induce intracellular signaling cascades. To understand where the receptors could potentially interact, we performed an in-silico protein-protein prediction in AlphaFold3 for direct contact between TLR4 and Siglec-5 (Figure S7). ^45^ For both Siglec-5 and TLR4, a per-residue pLDDT plot identified both confident (70+) secondary structures indicated by blue colors, as well as less confident structures (below 70) indicated by yellow and orange colors. Siglec-5’s intracellular domain consisted of intrinsically disordered regions (IDR), making predictions increasingly difficult. When analyzing the output data from the modeling, the predicted aligned error (PAE) in the relative position and orientations of the proteins revealed several potential interaction points between the proteins, both the resulting predicted template modeling (pTM) score of 0.47 and ipTM score of 0.21 indicated low confidence and an overall challenging prediction. However, the current limitations of glycan interactions in AlphaFold made it challenging to draw any valuable conclusions as Siglec-5 will bind to Sia.

We therefore used immunocytochemistry to investigate the spatial expression patterns and potential physical interactions on monocytes. Here we used an antibody with affinity to a protein sequence found in Siglec-5 and not Siglec-14. Representative images show localization of TLR4 (green) and Siglec-5 (red) under each condition (Figure 3A). Both Siglec-5 and TLR4 displayed uniform expression at 0hrs, which changed after 24hrs in culture. As with flow cytometry, a decrease was noted in TLR4 by LPS, whereas Siglec-5 maintained a uniform expression pattern on the cell membrane. At 0hrs, Mander’s colocalization coefficient (MCC), determined a strong TLR4 overlap with Siglec-5 (MCC = 0.78), while the more abundantly expressed Siglec-5, as expected, displayed a weak overlap to TLR4 (MCC = 0.35; Figure 3B and 3C). After 24hrs of stimulations, all conditions led to a reduction in TLR4 overlap with Siglec-5, with MCC values ranging between 0.46-0.50, indicating a moderate overlap. Cells stimulated with LPS had lost the correlation between Siglec-5 and TLR4 and now showed no evidence of interaction.

**Figure 3.**
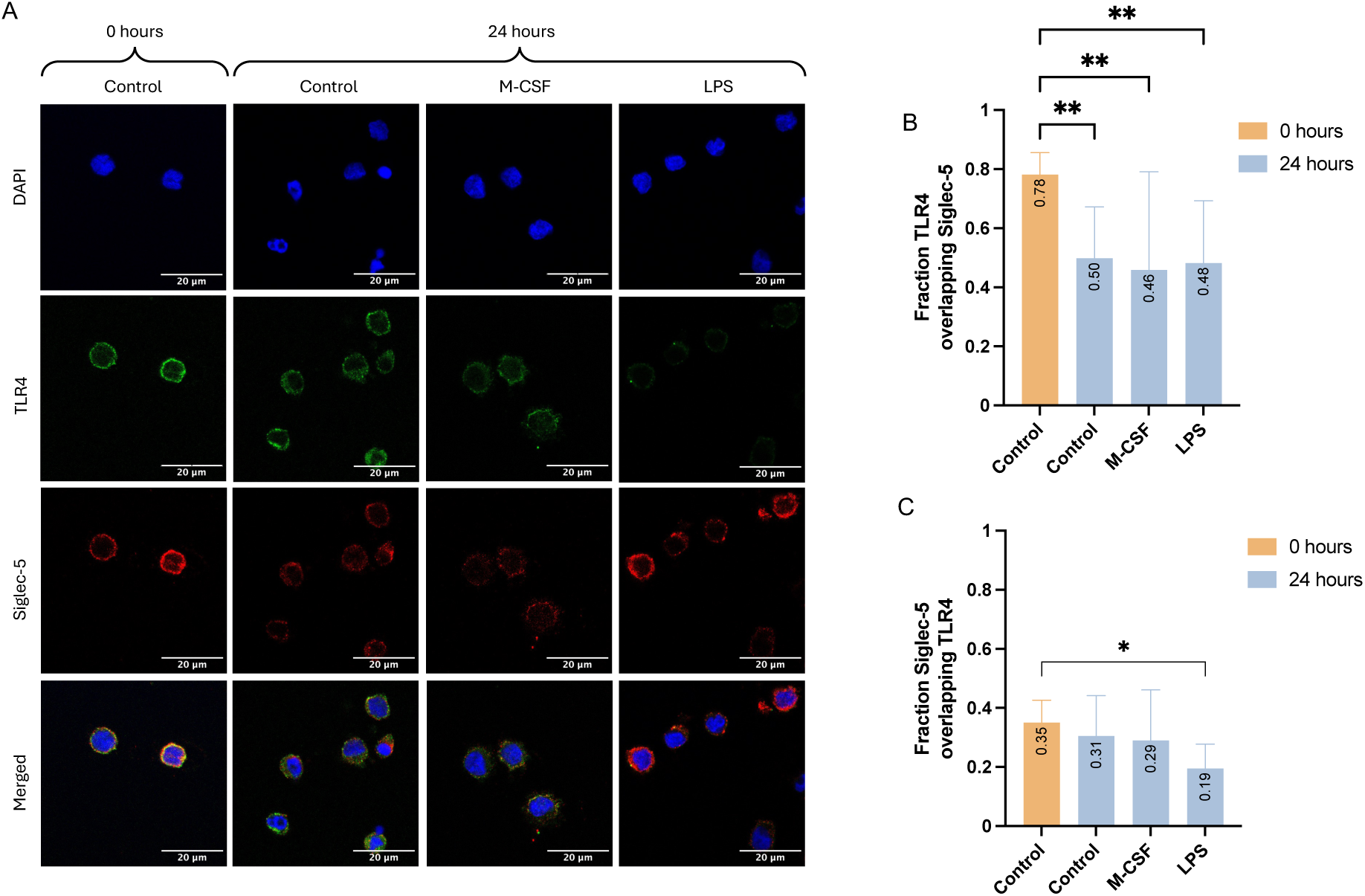
Siglec-5 and TLR4 co-localization on the cell surface of monocytes. (A) Representative immunofluorescence images of monocytes at baseline, and control, M-CSF, and LPS after 24hrs. Blue color (dapi) colored the nucleus, green color represents TLR4, and red color represents Siglec-5. Scale bar represents 20µm. (B-C) Mander’s colocalization coefficient (MCC) M1: Fraction TLR4 overlapping Siglec-5, and M2: Siglec-5 overlapping TLR4. Data is represented as mean±SD generated from 4-7 images per condition with two biological replicates. Statistical analysis was performed using one-way ANOVA with Dunnett’s multiple comparisons test (\**p*<0.05, \*\**p*<0.01).

In conclusion, the immunocytochemistry data indicated a potential for these receptors to physically interact on the cell membrane and be regulated by M-CSF and LPS.

### Intracellular inhibition of TLR4 reduced cytokine secretion and Siglec-5 expression

To assess the functional consequences of Siglec-5/14 and TLR4 interactions, we measured cytokines produced by the monocytes after the stimulations. To deduce if NFkB signaling was the main signaling pathway involved in the interactions we used the small molecule TLR4 inhibitor TAK-242. This inhibitor specifically binds to Cys747 in the intracellular domain of TLR4, inhibiting both MyD88-dependent and MyD88-independent pathways. ^46^ The involvement of TLR4-signaling was determined by evaluating how effective TAK-242 was in suppressing cytokine production using the bio-plex cytokine screening method. The inhibition was performed prior to the addition of M-CSF, IL1β/TNFɑ, and LPS and was maintained throughout the experiment.

LPS lead to the most pronounced increase in pro-inflammatory cytokines at both 4hrs (Figure S8A) and 24hrs (Figure 4A). With elevated levels of TNFɑ, RANTES, IL1β, IL-1ra, MCP-1, IL-10, and G-CSF detected, aligning with its known role to trigger TLR4-mediated inflammation. M-CSF, in contrast, was associated with lower cytokine secretion consistent with its role in differentiation of monocytes into macrophages. Prominent inhibition of cytokine secretion by TAK-242 was observed after LPS as well as with M-CSF stimulation after both 4 and 24 hours. IL1β/TNFɑ induced cytokine secretion as well, however the TAK-242 inhibitor had no influence on the signaling and levels of cytokines produced, as expected (Figure S9).

**Figure 4.**
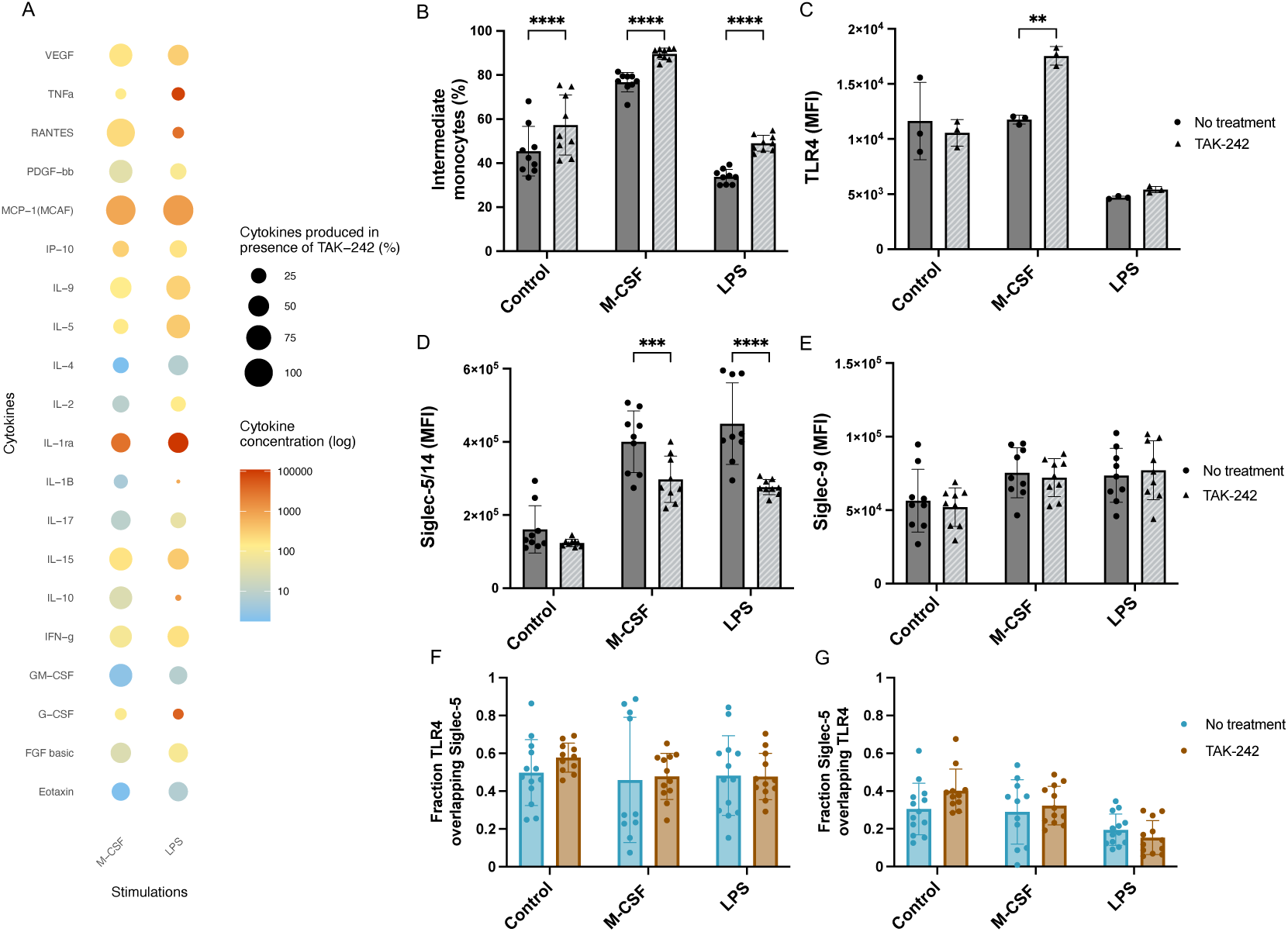
Intracellular inhibition of TLR4 reduced cytokine secretion and Siglec-5 expression. (A) Using the Bio-Plex Pro Human Cytokine Grp I Panel 27-Plex, a panel of cytokines and chemokines was screened in the supernatants of monocytes incubated for 24hrs with M-CSF and LPS, with and without the TLR4 inhibitor TAK-242. Each datapoint is the mean of three biological replicates with three technical replicates pooled together. Cytokine levels from monocytes stimulated in absence of TAK-242 is indicated by “Cytokine concentration (log)” and the “Cytokines produced in presence of TAK-242 (%)” indicates how effective TAK-242 was at limiting cytokine secretion compared to without TAK-242. (B) The percentage of intermediate monocytes after TAK-242 treatment was quantified in monocytes stimulated with M-CSF and LPS and compared to the control at 24hrs. Each datapoint represents one technical replicate. Three technical replicates were used from three biological replicates. (C-E) The effect of TAK-242 was assessed based on MFI of membrane-associated TLR4, Siglec-5/14, and Siglec-9 in monocytes stimulated with M-CSF and LPS and compared to the control at 24hrs. Each datapoint represents one technical replicate. Three technical replicates were used from three biological replicates, except for TLR4 which was analyzed from one biological replicate. (F-G) Mander’s colocalization coefficient (MCC) displaying fraction TLR4 overlapping Siglec-5 and vice versa. Each datapoint represents one image, 4-7 images were used per condition from two biological replicates. Data is represented as mean±SD, statistical analysis was performed using two-way ANOVA with Sidak’s multiple comparisons test (\**p*<0.05; \*\**p*<0.01; \*\*\**p*<0.001; \*\*\*\**p*<0.0001).

Next, we evaluated how blocking TLR4 affected monocyte subsets, including the intermediate population, as well as the receptors TLR4, Siglec-5/14, and Siglec-9. After 4hrs of stimulation, no significant changes in expression patterns were observed after TAK-242 treatment (Figure S8B-S8E). However, inhibition of TLR4 signaling induced a significant abundance in intermediate monocytes after 24hrs (Figure 4B). TAK-242 only induced significant TLR4 expression after M-CSF activation (Figure 4C). M-CSF and LPS induced the highest levels of Siglec-5/14 and interestingly, blocking TLR4 significantly reduced Siglec-5/14 expression in these two groups (Figure 4D). In contrast, Siglec-9 expression remained unchanged when TLR4 was blocked (Figure 4E). To determine whether TLR4 blockade affected both Siglec-5 and Siglec-14 similarly, we used the Siglec-14 specific antibody and found no significant change in surface expression (Figure S10). TAK-242 had no impact on the fraction TLR4 overlap with Siglec-5 and vice versa (Figure 4F and 4G). These findings suggest that TLR4 inhibition influence Siglec-5 expression.

### Siglec-5 and TLR4 interaction is abrogated by loss of sialoglycans

To delineate mechanisms pertaining to Siglec-5 signaling and its correlation to TLR4, sialidase was used to remove membrane-bound Sia and to assess its effect on cell behavior, especially as TLR4 is sialylated. ^38^ The ability of sialidase to enzymatically remove Sia from glycans was confirmed by LC-MS and lectin staining. An increased level of O-glycans lacking Sia was detected in sialidase treated monocytes (Figure 5A), as well as lower levels of ɑ2,3 and ɑ2,6 Sia in presence of sialidase (Figure 5B and 5C). TLR4 expression was also shown to be reduced after treatment with sialidase, whereas Siglec-5/14 and Siglec-14 displayed similar expression as control (Figure 5D and 5E, Figure S10). The colocalization of TLR4 and Siglec-5 after 24hrs of sialidase treatment was evaluated through immunocytochemistry, which revealed reduced expression of both receptors compared to the control at 0hrs (Figure 5F). The fraction overlap of TLR4 to Siglec-5 (M1), and Siglec-5 to TLR4 (M2) was evaluated using the MCC score (Figure 5G). This revealed a medium TLR4 overlap to Siglec-5 (MCC = 0.47). Similar to LPS, no evidence of overlap between Siglec-5 to TLR4 (MCC = 0.13) was observed. When measuring cytokine production using the bio-plex screening method, treatment with sialidase led to an increase in TNFɑ, RANTES, IL1β, IL-1ra, MCP-1, IL-10, and G-CSF, the same proinflammatory cytokines as LPS induced (Figure 5H).

**Figure 5.**
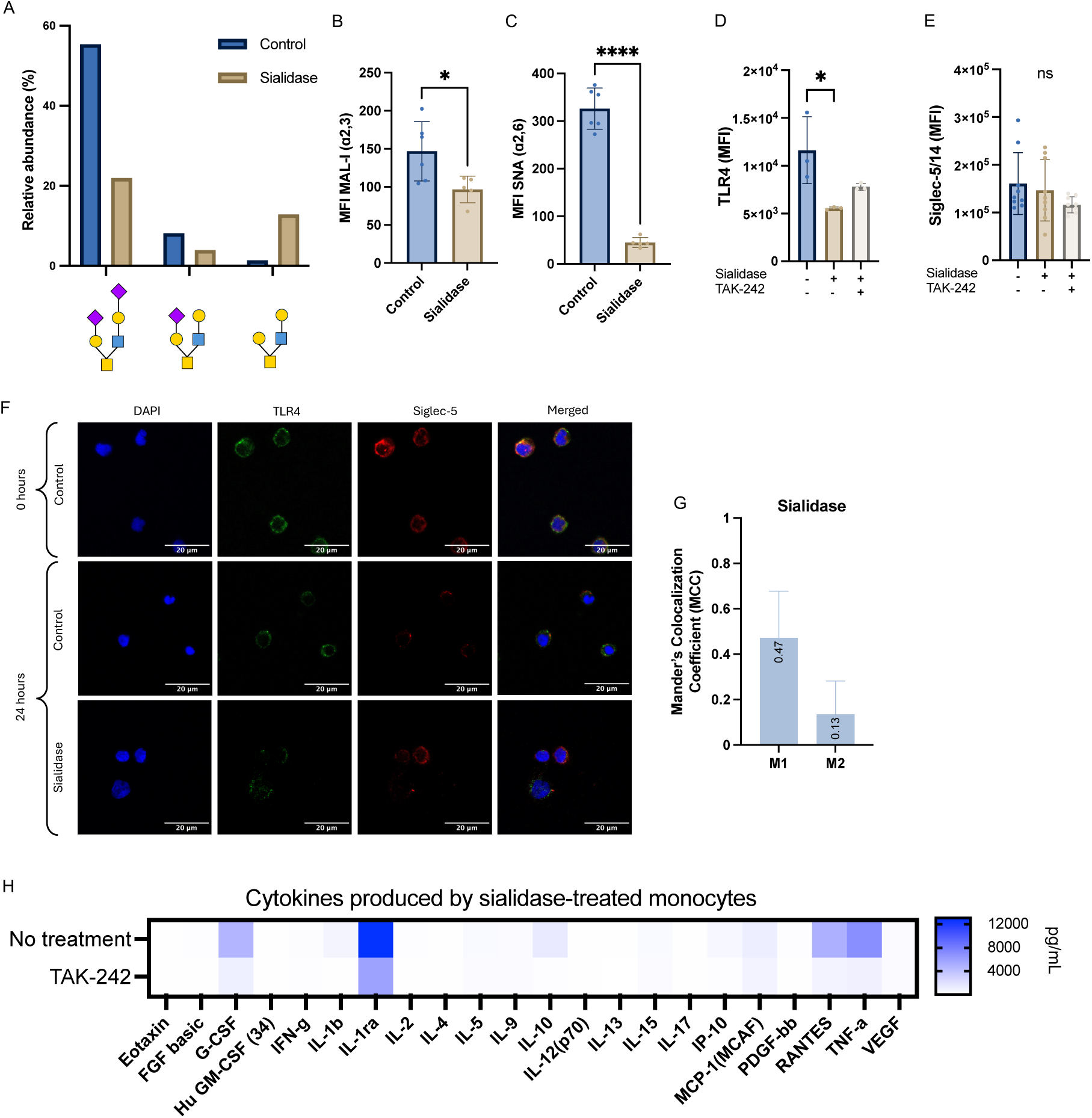
Siglec-5 and TLR4 interaction is abrogated by loss of sialoglycans. (A) Sialidase activity was confirmed using LC-MS, the relative abundance of top two sialylated *O*-glycans and non-sialylated *O*-glycan in control and sialidase treated monocytes was quantified. (B-C) ɑ2,3 and ɑ2,6 Sia on monocytes was quantified in three technical replicates from two biological replicates using MAL-I and SNA lectins. Both linkages significantly decreased in presence of sialidase after 24hrs in culture. (D-E) Monocytes were enzymatically stripped of their membrane-bound Sia for 24hrs in presence or absence of TAK-242 and compared to the control. MFI of membrane-associated TLR4, Siglec-5/14 and Siglec-14 was assessed in triplicates from one biological replicate for TLR4 and three for Siglec-5/14. (F) Colocalization of TLR4 and Siglec-5 was investigated in monocytes from two biological replicates treated with sialidase for 24hrs compared to the control at 0 and 24hrs. (G) Manders colocalization coefficient (MCC) M1 and M2, representing “Fraction TLR4 overlapping with Siglec-5” and “Fraction Siglec-5 overlapping with TLR4”, respectively. Values were acquired from 4-7 images per slide. (H) A panel of cytokines and chemokines was screened using the bio-plex assay in the supernatants of monocytes incubated for 24hrs with sialidase, with and without the TLR4 inhibitor TAK-242. Each datapoint is the mean of three biological replicates with three technical replicates pooled together. Data is represented as mean±SD, statistical analysis was performed using one-way ANOVA with Dunnett’s multiple comparisons test (\**p*<0.05; \*\**p*<0.01; \*\*\**p*<0.001; \*\*\*\**p*<0.0001).

Inhibition of TLR4 signaling using TAK-242 in cells treated with sialidase seemed to affect TLR4 surface expression but not Siglec-5/14. Similar to LPS, TAK-242 treatment of cells treated with sialidase was also effective in limiting cytokine secretion, suggesting that removing extracellular Sia on TLR4 induces both dependent- and independent MyD88 pathways. ^46^

### IL-6 secretion levels can be linked to distinct monocyte phenotypes

The secretion of TLR4-dependent IL-6 production induced by sialidase, LPS, and M-CSF, was compared to monocytes stimulated with OA SFs using a sandwich ELISA. Sialidase and LPS induced the highest levels of IL-6 and OA SFs induced a low-grade inflammatory profile (Figure 6A).

**Figure 6.**
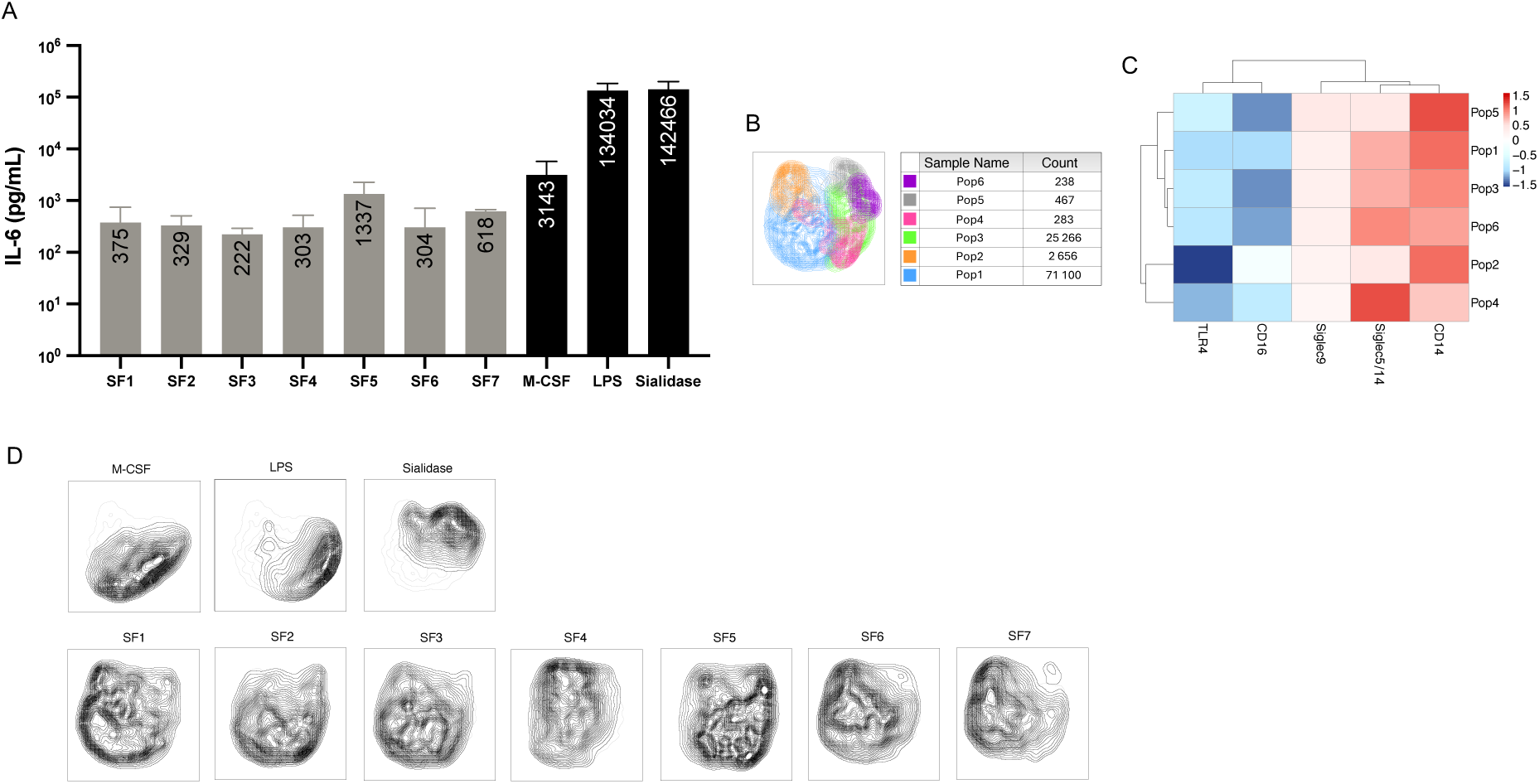
IL-6 secretion levels can be linked to distinct monocyte phenotypes. (A) The concentration of IL-6 (pg/mL) produced by the stimulated monocytes were quantified using a sandwich ELISA, grey bars represent SFs, and black bars represent the pathway-specific stimulations. Data is represented as mean±SD, with three technical replicates from three biological replicates. (B) Analysis was performed on concatenated monocytes stimulated with M-CSF, LPS, sialidase, and SF1-7. Subsequently, monocytes were downsampled to 100,000 cells before performing clustering analysis using the FlowSOM plugin. This generated six monocyte subpopulations, the table displays each population and its count, respectively. For the monocytes stimulated with SF1-7, three technical replicates were used from three biological replicates. For monocytes stimulated with M-CSF, LPS, and sialidase, three technical replicates were used from one biological replicate. (C) Heatmap analysis demonstrate the six populations and their relative expression of TLR4, CD16, Siglec-9, Siglec-5/14, and CD14. (D) Each UMAP contour density plot represents one condition, i.e. M-CSF, LPS, sialidase, and SF1-7.

By understanding how the patient IL-6 profiles corresponded to the phenotypes induced from the Siglec-5 – TLR4 interactions, clustering analysis was performed based on CD14, CD16, TLR4, Siglec-5, and Siglec-9. Clustering was performed on three biological replicates from monocytes stimulated with OA SF, whereas one biological replicate was used from the stimulations. All groups were downsampled to view 10.000 cells. Uniform Manifold Approximation and Projection (UMAP) dimensionality reduction (Figure 6B) displayed the six distinct monocyte populations generated (Pop1-Pop6), with Pop1 being the largest (71,100 cells) and Pop6 being the smallest (238 cells). Heatmap analysis of marker expression (Figure 6C) revealed that Pop1 was characterized by reduced TLR4/CD16 and increased Siglec-5/14/CD14 expression. Pop2 displayed the lowest levels of TLR4, accompanied by an induced CD14 expression. Siglec-5/14 was upregulated in Pop1, 3, 4, and 6. Each condition was displayed as UMAP density contour plots (Figure 6D). Pop1 and Pop4, along with Pop2 were the most abundant subtypes generated in monocytes stimulated with SF1-7, revealing similarities with the low-grade inflammatory profile induced by M-CSF. While monocytes stimulated with SFs displayed a wide distribution of subpopulations, it was evident that SF5 revealed similarities to the higher inflammatory conditions (LPS and sialidase) with a higher abundance of Pop3, Pop5, and Pop6 as well as a higher level of IL-6. These populations displayed the lowest expression of CD16, higher TLR4/CD14, and varying Siglec-5/14 levels.

## DISCUSSION

In this study, we identified Siglec-5/14 as a potential regulatory protein in OA pathogenesis using a unique approach involving NPs. By performing proteomics analysis on the proteins formed around NPs incubated with OA SF, we detected Siglec-5/14 as one of 14 most abundant proteins. Through the IntAct Molecular Interaction Database, TLR4 was identified as the protein with the highest confidence score to Siglec-5 but not to Siglec-14. While the role of TLR4 has been extensively studied in OA, ^22^ the link to Siglecs remain unclear. We confirmed the regulatory role of Siglec-5/14 on monocytes by stimulating cells with OA SF, inducing Siglec-5/14 expression which inversely correlated with TLR4. Mechanistically, we showed that the potential Siglec-5 – TLR4 interactions are regulated by sialoglycans on the cell surface, where highly inflammatory cytokines such as those induced by LPS or sialidase, closely mirrored those found in patients with high inflammatory activity. Since TLR4 is increasingly recognized as a key player in age-related diseases like OA, ^22,23,27,47^ our finding that Siglec-5 can modulate TLR4-driven proinflammatory signaling highlights a potentially unexplored regulatory axis in OA pathogenesis.

Interestingly, we found that OA SF induced Siglec-5/14 expression which inversely correlated with TLR4 at 24hrs, suggesting potential crosstalk between these receptors. This inverse relationship may reflect a negative feedback mechanism, where Siglec-5 reduces TLR4-driven inflammation in OA, potentially mitigating tissue damage. To deduce Siglec-5-mediated regulation of TLR4, SHP-1 was inhibited using TPI-1, and monocytes were stimulated with LPS for 4hrs. Inhibition of SHP-1 resulted in an increased production of key TLR4-associated cytokines, including IL-6, IL-8, and MCP-1, while levels of IL1β and TNFɑ either decreased or remained the same. While these findings present some conflicting patterns, they suggests Siglec-5 contributes to TLR4 regulation, with potentially more pronounced effects at later timepoints. Siglec-5 has been shown to interact with TLR4, and the murine Siglec-E facilitates TLR4 internalization. ^21,32,48^ Given that TLR4 is a key driver of proinflammatory activity in OA, ^22,47^ its regulation by Siglec-5 may serve as an important immunomodulatory checkpoint.

This was further strengthened by the observation that LPS and sialidase significantly reduced TLR4 expression, likely as a desensitization mechanism to prevent monocytes from overreacting to sustained inflammatory stimuli. ^49^ We investigated potential direct interactions between the two receptors, as TLR4 is known to be heavily sialylated. ^38,50^ Using a Siglec-5 specific antibody, colocalization analysis revealed a strong overlap at baseline, in line with previous studies suggesting that Siglecs can bind and regulate TLR activity. The high level of IDRs at the membrane interface in both TLR4 and Siglec-5 indicate a tunable interaction surface, one capable of rapid regulation in response to environmental cues, a key feature of immune cell communication. ^51^

However, with prolonged stimulations, we observed a progressive reduction in colocalization. Colocalization scores below 0.2 indicated a loss of interaction, suggesting that LPS and sialidase disrupted Siglec-5/TLR4 engagement. This supports a model where the interaction is temporally regulated, strong early on then declining as immune activation progresses. This temporal tuning may serve as a critical checkpoint, allowing for transient modulation of TLR4 signaling during early immune responses while ensuring desensitization under persistent stimulation to prevent tissue damage.

Mechanistically, the loss of interaction appears to stem from depletion of membrane-bound Sia rather than complete receptor downregulation. Surface Sia levels have been directly linked to a cells’ maturation state and inflammatory potential, ^52,53^ and their removal, via sialidase or inflammatory stimuli, may directly influence Siglec binding. Both Siglec and TLR4 expression and activity are modulated by interactions with glycan structures, particularly ɑ2,3 and ɑ2,6 Sia linked residues. These glycan modifications can modulate immune cell signaling and are often altered in response to inflammation ^54,55^.

The role of Siglec-5 in OA and its exact signaling mechanisms remain poorly understood. Our study found that short-term activation reduced Siglec-5/14 expression, which was recovered by 24hrs, suggesting either a shedding or internalization mechanism. A recent study on septic patients found elevated levels of soluble Siglec-5 in plasma which could be induced by stimulating monocytes with LPS. ^17^ Monocytes might hence shed Siglec-5 to modulate other immune cells, explaining why Siglec-5 could be found in high abundance in the proteins associated with the NPs. It is possible that Siglec-5 plays a role in this desensitization to not exacerbate inflammation. Siglec-14, which has an opposing immunological role to Siglec-5, has also been detected as a soluble protein, ^20^ however, the absence of changes in membrane expression across different stimulations-when assessed using a Siglec-14-specific antibody, suggests that the effect observed with the Siglec-5/14 targeting clone 1A5 are predominantly driven by Siglec-5. Monocytes stimulated with LPS and sialidase exhibited highly similar cytokine secretion profiles, reflecting their shared activation through TLR4. ^37,38,56^ Other studies have shown that LPS leads to an overexpression of host sialidase, reducing membrane-bound Sia, and using an exogenous sialidase can thereby mimic LPS stimulation. ^57^ This functional similarity was further supported by the comparable inhibitory effects observed by TAK-242. Several studies ^36,58^ have demonstrated changes in surface sialylation in OA chondrocytes and fibroblast-like synoviocytes in rheumatoid arthritis, indicating that inflammation significantly affects glycosylation patterns. ^36,58^ Our UMAP analysis revealed a wide distribution of both low- and high-grade inflammatory monocytes, with proinflammatory profiles generated by LPS and sialidase. Interestingly, SF5 exhibited an overlap with a high-grade inflammatory profile, correlating with increased IL-6 secretion.

Other OA SF-stimulated monocytes shared similarities with M-CSF-stimulated monocytes, consistent with a low-grade inflammation. These findings highlight how interpatient variability influence subtle changes in monocyte surface markers, ultimately shaping inflammatory responses, where Siglecs could play a role in regulation. The high levels of Siglec-5 induced by LPS and contrasting low levels induced by sialidase, indicated that the immunomodulatory effects by Siglec-5 was limited in high-inflammatory conditions within 24hrs. However, in low-grade inflammation, such as with OA SF, patients with higher Siglec-5 expression may have a more tightly controlled immune response, preventing the overactivation of TLR4 signaling and associated inflammation.

In conclusion, this study highlights the role of Siglec-5 on TLR4 in the pathogenesis of OA, shedding light on their intricate interplay. While both Siglec-5 and TLR4 are implicated in the inflammatory process of OA, their complex relationship warrants further investigation to fully elucidate the underlying mechanisms and interactions and the contributing effects by Siglec-14. The dynamic interplay between glycosylation, receptor localization, and immune signaling could in the future be validated using proximity ligation assays, live cell imaging or co-immunoprecipitation to further clarify the spatial and temporal nature of this interaction, while lectin-based profiling could confirm changes in Sia presentation. The marked reduction in cytokine production upon TLR4 inhibition underscores the therapeutic potential of TAK-242 particularly for patient subgroups with low Siglec-5 or skewed Sia expression, where modulation of TLR4 signaling may prove especially beneficial. Similar features imposed on monocytes by LPS and sialidase demonstrated TLR4-specific activation. Yet, sialidase failed to upregulate Siglec-5, emphasizing an important role of Sia as ligand which could serve as a potential therapeutic target for immune cell modulation. Restoring Siglec-5 function and understanding its signaling mechanisms presents promising opportunities to modulate TLR4 signaling and reduce inflammation not only in OA, but for many pathogeneses where modulating glycosylation patterns can be used to fine-tune immune responses.

## Limitations of the study

We acknowledge some key limitations to our study. Employing an antibody that specifically recognizes Siglec-5 would help avoid ambiguity arising from cross-reactivity with Siglec-14, as is possible with the 1A5 clone. While this study has focused on healthy monocytes, many different cells are involved in OA, including chondrocytes and synoviocytes. Modulation of CD8^+^ T-cell proliferation by soluble Siglec-5, ^17^ suggests Siglec-5 could have potential effects on other OA-related cells. The transient increases in Siglec-5, alongside persistent decreases in the sialidase group, suggest ongoing glycan remodeling that may involve TLR4 or other receptors. Longer observation periods and dose-dependent studies, particularly with sialidase, are needed to clarify these dynamics. Broader investigation into glycosylation patterns and their impact on immune receptor networks could uncover additional therapeutic targets. Future work should also validate these findings in vivo and in human OA samples, particularly OA-derived monocytes, focusing on how sialylation patterns and Siglec-5 expression vary among patient subgroups.

## Resource availability

### Lead contact

Further information and requests for resources should be directed to and will be fulfilled by the lead contact, Alexandra Stubelius.

### Material availability

The study did not generate new materials.

### Data and code availability

- All data can be obtained from the lead contact, provided the request is reasonable.
- The code related to the algorithm can be accessed by reaching out to the lead contact.
- Any additional information required to reanalyze the data reported in this paper is available from the lead contact upon request.

## Acknowledgements

Anna-Karin Hultgård-Ekwall, Lena Björkman, Thomas Eisler, and Ola Rolfson are gratefully acknowledged for arranging the patient samples, and Sofia Grindberg and Paula-Therese Kelly Pettersson at Danderyd’s Hospital and Lotta Falkendahl at the University of Gothenburg are acknowledged for their assistance in collecting the samples. This work was supported by Chalmers Technical University and its Area of Advance Nano. The authors greatly acknowledge further financial support from the Foundation for Sigurd and Elsa Goljes Minne (LA2021-0100), The Royal Swedish Academy of Sciences (CR2021-0024), The Jeanssons Foundation (J2021-0050), The Hasselblad Foundation, the Swedish Research Council (2021-01870), Wilhelm och Martina Lundgrens Vetenskapsfond (2022-4054) the Swedish Rheumatism Association (R-981253), the King Gustav V’s 80 years’ foundation (FAI-2022-0872) and IngaBritt och Arne Lundbergs Forskningsstiftelse (LU2022-0041).

## Author Contributions

Study design (L.R, A.S), conducting experiments (L.R, F.J, U.v.M, V.V), acquiring data (L.R, F.J, U.v.M, V.V), analyzing data (L.R, F.J, U.v.M, V.V, A.S), providing funding and reagents (N.K, A,S), writing the manuscript (L.R, A.S). All authors have contributed to editing and approved the manuscript.

## Declaration of generative AI and AI-assisted technologies in the writing process

During the preparation of this work, we used OpenAI for language editing and to ensure readability. After using this tool/service, we reviewed and edited the content and take full responsibility for the content of the published article.

## STAR METHODS

### EXPERIMENTAL MODEL AND STUDY PARTICIPANT DETAILS

#### Sex as a biological variable

This study includes patient material from both female and male patients. Synovial fluid from 7 OA patients (2 females) due for total knee replacement surgery was selected for the experiments: age 63-87 (Median 77) and BMI 23.4-38.0 (Median 25.6). Due to the small patient population, no further stratification was made. To remove cells from OA SF, the SFs were centrifuged at 3000 x g for 10 min before being stored at -80°C.

#### Primary cultures

Buffy coats were obtained from anonymous healthy donors from the blood bank at Sahlgrenska University Hospital, Gothenburg, Sweden (18/23). The sex of the cells could not be reported as they were obtained from anonymous donors. Isolated cells were cultured at 37°C with 5% CO_2_.

## METHOD DETAILS

### Protein corona isolation and material synthesis

NP synthesis was previously described in von Mentzer *et al.* ^10^ Briefly, PAMAM (G5) were modified by PEGylation using PEG 350 MW, NP_350_ at 2%, or PEG 5000 MW, NP_5000_ at 2%, purified, and used at the final concentration of 30 μM in PBS (pH = 7.4).

SF samples from 4 (2f/2m, 45–82 years old) late OA patients were collected during aseptic aspiration of knee joints at the Rheumatology Clinic and Orthopaedic Clinic, respectively, at Sahlgrenska University Hospital, Gothenburg, Sweden.

Patient SF were pooled, diluted 1:20 in PBS, and mixed with 30 μM NP solution (1:1, v/v). The samples were incubated at 37°C while shaking for 1 h at 100 rpm to resemble the dynamic environment of the SF in the joint and address the Vroman effect. To preserve the proteins bound with high affinity known as the hard protein coronas, NPs were spun down at 15,000 × *g* for 15 min and washed three times with chilled PBS. Final concentration of NP-protein complexes was 1 μg/mL (2.5 × 10^8^ particles/mL).

NPs were pelleted by centrifugation at 15,000 × *g* for 15 min, snap-frozen using liquid nitrogen, and submitted to the Proteomics Core Facility (Gothenburg, Sweden). Briefly, proteins were digested into peptides using MS-grade trypsin and analyzed by nanoscale liquid chromatography-tandem mass spectrometry LC-MS/MS. The mascot search engine was used to match the discovered peptide sequences against SwissProt human and bovine protein database using Proteome Discoverer. Data were analyzed using a label-free quantification method and the protein false discovery rate was set to 1%. To elucidate the molecular functions and classifications of significant proteins, the enrichment analysis was performed using Gene Ontology-based PANTHER classification system. SF samples were matched to human (Homo sapiens). The detected proteins were analyzed and illustrated as a volcano plot using the DEP R package as described in. ^10^

### Monocyte isolation

Buffy coats were prepared at Sahlgrenska hospital the day before and collected for the experimental study in the morning. Monocyte isolation was performed using the negative selection RosettSep^TM^ Human Monocyte Enrichment Cocktail. Since this kit is optimized for whole blood, the buffy coat was diluted 1:1 with Dulbecco’s phosphate buffered saline (PBS) (Gibco) to ensure that the concentration of nucleated cells did not exceed the limit. Apart from that, the isolation was performed according to the protocol. The blood:PBS was mixed with the cocktail and 1mM EDTA and incubated for 20 min at room temperature. Following this, the mixture was placed over a Lymphoprep™ density gradient medium and centrifuged in a swing-out bucket for 20 min at 1200 x g with the brake off. The enriched monocytes were transferred to a new tube and washed in the recommended medium containing PBS, 2% heat inactivated fetal bovine serum (FBS) and 1mM EDTA. The monocytes were washed three times in recommended medium and centrifuged at 300 x g for 10 min with the brake low for the first two washes. For the final wash step, the cells were centrifuged for 120 x g for 10 min with the brake off to remove platelets.

### Monocyte stimulations

The enriched monocytes were counted and plated at a concentration of 3 × 10^5^ cells/well in a 48 well plate. The cells were diluted in RPMI Medium supplemented with 1 % glutaMAX, 1 % Penicillin/Streptomycin and 10 % FBS. TAK-242 or TPI-1 was added to the designated wells at a concentration of 1µM or 100ng/mL, respectively, and incubated for one hour. The final concentration of each stimulation was 10ng/mL of IL1β and TNFɑ, 100ng/mL M-CSF, 100ng/mL LPS, and 100mU/mL sialidase. The total volume per well was 250µL. The cells were analyzed after 0, 4, and 24 hours.

As a comparison to these commonly used stimulations, we incubated primary monocytes with OA synovial fluid at a concentration of 25%. After 24hrs of stimulation the surface markers were detected using flow cytometry.

### Cell staining for flow cytometric analysis

Monocytes were collected by transferring the whole volume to a v-plate and centrifuging them at 300 x g for 5 minutes, 4°C. The supernatants were transferred and saved at -80°C for cytokine measurement. Meanwhile, to detach the monocytes that started differentiating into macrophages, 50µL trypsin-EDTA was added. Once detached, 50µL ice cold recommended medium was used to inactivate trypsin. The detached cells were added to the v-plate and washed twice. 10µL Fc block was added to each well and the plate was incubated on ice for 10 min. Monocyte phenotypes were analyzed using the following antibodies at a final concentration of 1:100, CD14-FITC, CD16-Brilliant Violet 605, Siglec-5-PE, Siglect-9-PE-Cy7, and TLR4-APC. 10µL the antibody cocktail was added to each well and the plate was incubated for 20 min on ice in the dark. UltraComp eBeads^TM^ Plus Compensation Beads were used to compensate spillover. The cells and compensation beads were washed three times in recommended medium. To exclude dead cells, 3µM DAPI was added. Fluorescence minus one (FMO) and unstained cells were used to set up the gating for each individual experiment. The cells were run on the Cytoflex LX (Beckman Coulter), median fluorescence intensity (MFI) was recorded and analyzed using FlowJo (Version 10.10.0). ^59^

### Unsupervised flow cytometric analysis

Unsupervised analysis was performed using FlowJo. Cells from all stimulations (SF1-SF7 and M-CSF, LPS, and sialidase) were concatenated into one single file. Clustering was conducted using the FlowSOM plugin, generating six distinct populations. Subsequently, a UMAP was generated based on these populations. For visualization, each stimulation condition was overlaid onto separate UMAPs to illustrate the distribution of populations across each stimulation.

### LC-MS

After isolating monocytes, they were diluted to a concentration of 2 × 10^6^ cells and added to two Eppendorf tubes. Sialidase was added so that the number of units per cell was the same as in the experiments, the tubes were incubated for 1 hour at 37°C. The tubes were centrifuged at 2500 x g for 10 min at 4°C. Supernatant was removed, the same washing step was performed two more times. Subsequently, 100µL lysis buffer (PBS + 2% SDS) was added, the tubes were briefly sonicated followed by gentle shaking for 10 min. Cell debris was removed by centrifugation at 14000 x g for 20 min. The supernatant was collected, and the protein concentration was measured using gold-standard BCA assay. Approximately 30 µg of extracted protein samples were dot blotted on a PVDF membrane (Millipore). Proteins were stained using Alcian blue stain solution, excised and transferred to an Eppendorf tube.

O-glycans were released and analyzed on liquid chromatography and mass spectrometry according to this protocol. In short, the dot blotted proteins were subjected to reductive β-elimination with 0.5 M sodium borohydride in 50 mM sodium hydroxide for 16 h at 50°C to release the glycans. The reduction reaction was quenched by the addition of glacial acetic acid and the resulting glycan containing mixtures were desalted using strong cation exchange resin packed on top of C18 column to remove the remaining proteins or peptides. The released glycans were analyzed using liquid chromatography and mass spectrometry. The glycans were separated on a column (10 cm × 250 µm) packed in-house with 5 µm porous graphite particles (PGC, Hypercarb, Thermo-Hypersil, Runcorn, UK) and at a flow rate of 6 μl/min. The samples were analyzed in negative ion mode on an Orbitrap mass spectrometer (Fusion, Thermo Electron, San José, CA) and the data acquisition was conducted with Xcalibur software (Version 2.0.7).

### Bio-Plex Pro Human Cytokine Panel 27-Plex

Supernatants from the monocyte stimulation assays were screened for a panel of 27 cytokines and chemokines to evaluate the immune response in the different conditions. The technical replicates from three biological replicates were combined, respectively, to reduce the number of wells to one well per biological replicate. The assay was performed according to the manufacturer’s instructions. Vortexed beads were added to each well, followed by two washing steps. The supernatants were centrifuged at 1,000 x g for 15 min at 4°C prior being added to the plate along with the standard, blank, and control. The plate was incubated shaking at 850 rpm for 30 min, followed by three washing steps. Detection antibodies were added to each well and incubated shaking for 30 min. After three washings, streptavidin-PE was added and incubated shaking for 10 min. An additional three washings were performed before resuspending the beads in assay buffer and reading the plate.

For values that were out of range, extrapolated values were assumed based on the highest and lowest value of the standard curve. We observed that most of the groups was below detection level for IL-7 and above detection level for MIP-1ɑ, MIP-1β, IL-6, and IL-8 and they were hence removed. Additionally, the media control was below detection level for most analytes and thus removed.

The cytokines detected were normalized in Prism (GraphPad software Version 10.4.1 (532)). For each subcolumn, values were scaled as fractions, with 0% representing smallest value of each sub column, and 100% represents the sum of all values in the subcolumn. The ggplot2 R^60^ package was used to create the dot plot heatmap illustrating cytokine expression under different stimulation conditions, along with the percentage cytokines remaining in presence of TAK-242.

### IL-6 ELISA

To determine the quantity of IL-6 in the supernatants of monocytes stimulated with SFs and M-CSF, LPS, and sialidase, a sandwich ELISA was performed according to the manufacturer’s protocol. Briefly, the ELISA plate was coated with the capture antibody the day before and incubated overnight the fridge. In the morning the next day, the plate was washed four times and blocked for 1 hour, shaking at 500rpm. Meanwhile, the standard and sample dilutions were prepared and subsequently added after an additional four washes. The plate was incubated for 2 hours, shaking, then washed four times. Detection antibody was added, and the plate was incubated for 1 hour, shaking. The same washing procedure was performed and avidin-HRP was added and incubated for 30 min, shaking. After this, the plate was washed 5 times and TMB substrate solution (A+B) was added and incubated for 15 min in the dark. Stop solution was added and the absorbance at 450nm and 570nm was detected.

### LEGENDplex Human Inflammation Panel 1 (13-plex)

A panel of 13 cytokines were screened in the supernatants of monocytes stimulated for 4hrs with and without the TPI-1 inhibitor. Following the manufacturer’s instructions, assay buffer, sample, standard, and premixed beads were added to a v-bottom plate and incubated for 2hrs shaking at room temperature at 800 rpm in the dark. Two washed was performed by centrifuging at 250 x g for 5 min. The supernatant was removed by flicking the plate. Detection antibodies was added and the plate was incubated for 1hrs, at room temperature at 800rpm, in the dark. SA-PE was added and the plate incubated for an additional 30 min. Following this, the plate was washed twice and the beads were resuspended in wash buffer and detected on the Cytoflex LX (Beckman Coulter) and analyzed on Biolegends LEGENDplex software.

### Confocal microscopy

To analyze TLR4 and Siglec-5 colocalization, confocal imaging was performed. 1.2×10^6^ isolated monocytes were seeded in complete RPMI in 12 well plates and pre-treated with 1µM TAK-242 or PBS, for 1 hour. Following this, the stimulations M-CSF, LPS, and Sialidase were added accorded to previously mentioned concentrations. After 0 and 24h, the monocytes were transferred to Eppendorf tubes and the leftover attached cells were trypsinized and transferred to the tubes before being centrifuged at 1,500 rpm for 5 min. The supernatant was discarded, and the cells were washed with PBS and centrifuged to remove the supernatant, this was performed three times. To fix the cells, 100µL 4% paraformaldehyde was added to each tube and incubated at room temperature for 10 min. The monocytes were blocked for 30 min with 5% FBS. Detection was performed using anti-human TLR4 mouse monoclonal antibody and anti-human Siglec-5 rabbit polyclonal antibody, both diluted 1:100 and stained for 1h in room temperature. 2 µg/mL of secondary antibody goat anti-mouse IgG and goat anti-rabbit IgG was incubated in the dark for 1h. After washing, the cells were spun onto a glass slide, ProLong^TM^ Gold Antifade Mountant was added before mounting the cover slip.

To verify sialidase enzyme efficiency at 24hrs in culture, 1.2×10^6^ isolated monocytes were seeded in complete RPMI in 12 well plates with or without the enzyme. The fixation protocol was the same as above, with a few exceptions; blocking was performed for 30 min using carbo-free buffer. For detecting Sia, lectin staining was performed using 10µg/mL Maackia Amurensis-I (MAL-1)-FITC and 10µg/mL Sambucus Nigra (SNA)-Cy5, targeting ɑ2,3 and ɑ2,6 Sia, respectively.

Images were acquired by using an inverted Nikon C2+ confocal microscope (Nikon Instruments, Amsterdam, Netherlands) equipped with a C2-DUVB GaAsP detector unit offering variable emission bandpass settings. A 60x Nikon APO oil-immersion objective (numerical aperture 1.4). Excitation and emission were set as follows for different staining: DAPI - λ_ex_ 401 nm, λ_em_ 425-497 nm, goat anti-rabbit IgG (Alexa Fluor 647) - λ_ex_ 639 nm, λ_em_ 644-695 nm, goat anti-mouse IgG (Alexa Fluor 488) - λ_ex_ 489 nm, λ_em_ 503-546 nm, MAL-I (FITC) - λ_ex_ 489 nm, λ_em_ 513-544 nm, and SNA (Cy5) - λ_ex_ 639 nm, λ_em_ 646-689 nm.

### Image analysis

The confocal images displayed in Figure 4A were enhanced for illustrative purposes. The brightness was equally enhanced for TLR4 and Siglec-5, for accurate comparison. Mander’s colocalization coefficient (MMC) was performed on unaltered images. Threshold was set to 90 for TLR4 and 140 for Siglec-5 to remove background. Antibody validation was performed to evaluate cross-species binding, these controls demonstrated 0 overlap. MCC was calculated through the JACoP plugin in ImageJ2 (Version 2.16.0/1.54p). ^61^

Quantification of ɑ2,3 and ɑ2,6 Sia was performed by extracting the MFI from the original intensity profiles with no alterations.

### Protein prediction

The protein prediction was analyzed and saved using ChimeraX. ^62^

### R packages

Figure 1B, Figure 4A, Figure 6C, and Figure S7A were created in Rstudio (R version 4.3.2 (2023-10-31). The following packages were used: readr, tidyverse, igraph, ggraph, ggplot2, and scales.^63–69^

### Statistics

Statistical significance was performed using Prism (GraphPad software Version 10.4.1 (532)). Data are shown as the mean and standard deviation and statistical differences were analyzed using One-way ANOVA followed by Tukey’s or Dunnett’s multiple comparisons test or two-way ANOVA followed by Sidak’s multiple comparisons test. Correlation was analyzed using Spearman correlation analysis. All statistical details can be obtained in the figure legends.

### Study approval

The studies involving human participants were reviewed and approved by the Regional Ethical Review Board in Gothenburg. All patients provided informed consent, and the procedure was approved by the Ethics Committee of Gothenburg University (Ethical approval Dnr: 573-07, 172-15).

