## Supplemental information for "Sialoglycans Modulate Siglec-5 – TLR4 Interactions in Osteoarthritis"

**Affiliations:**

### SUPPLEMENTAL INFORMATION

**Figures S1-S8.**

**Figure S1: Gating strategy.**

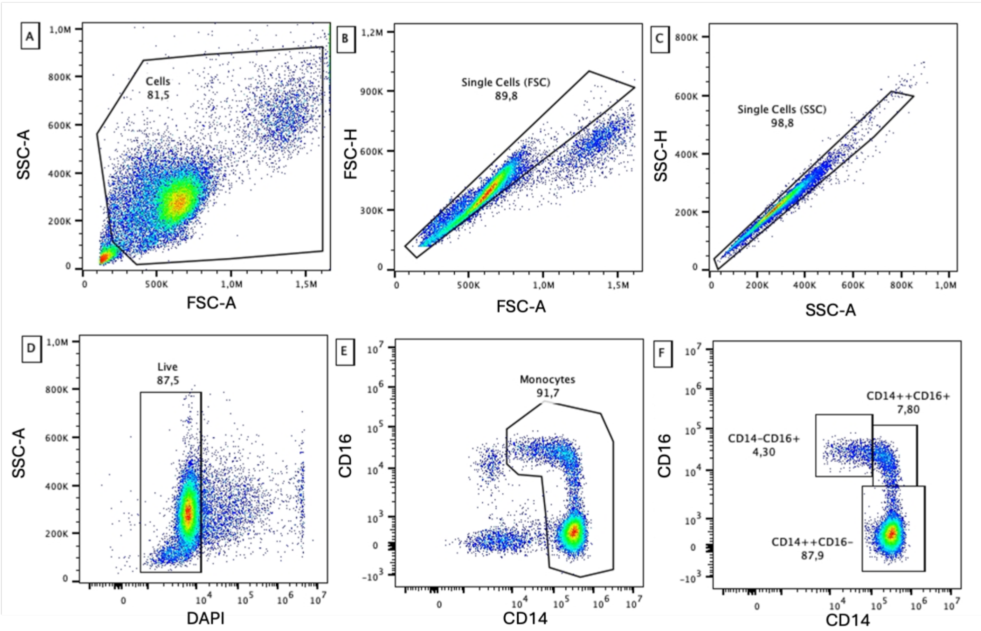


**Figure S1:** After isolating blood monocytes through negative selection, the cells were analyzed using flow cytometry. Debris was gated out using forward scatter area (FSC-A) versus side scatter area (SSC-A). Single cells were excluded first by putting FSC-A against FSC-H, followed by SSC-A against SSC-H. Live cells were detected by gating DAPI negative cells against SSC-A. Monocytic cells were gated out from the live cell population based on CD14 and CD16 expression, and non-monocytic cells were excluded. The ration between classical (CD14$^{+}$CD16$^{-}$), intermediate (CD14$^{+}$CD16$^{+}$), and non-classical (CD14$^{dim}$CD16$^{+}$) was determined.

**Figure S2: Monocyte isolation efficiency, percentage monocyte subsets, and expression of surface markers.**

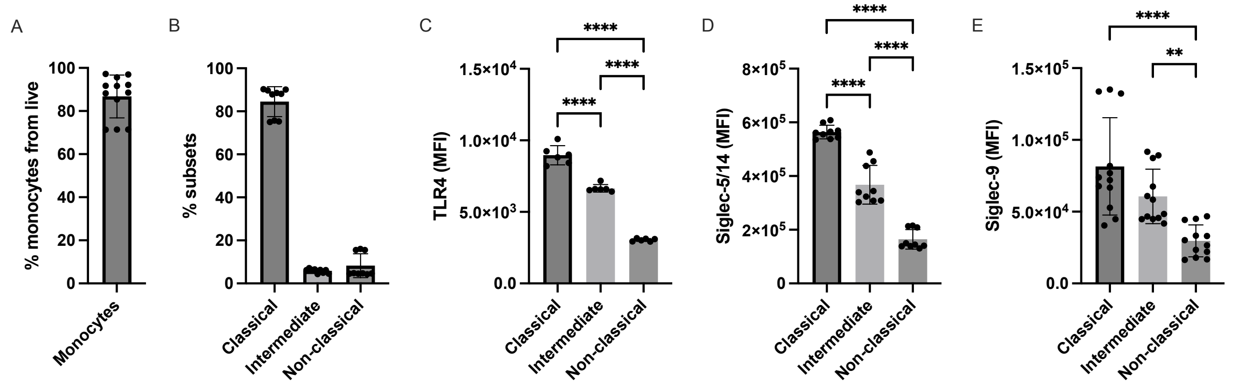


**Figure S2:** (A) The enriched monocyte fraction from the live cell population was confirmed to be between 72-85%, with a mean of 85.86%. (B) Percentage monocyte subsets at 0hrs. (C-E) Median fluorescence intensity (MFI) of TLR4, Siglec-5/14, and Siglec-9 in each monocyte subset at 0hrs. Data is represented as mean±SD with at least three technical replicates from three biological replicates. Statistical analysis was performed through one-way ANOVA using Dunnett’s multiple comparisons test (*$p$<0.05; **$p$<0.01; ***$p$<0.001; ****$p$<0.0001).

**Figure S3: Siglec-15 is not expressed by monocytes.**

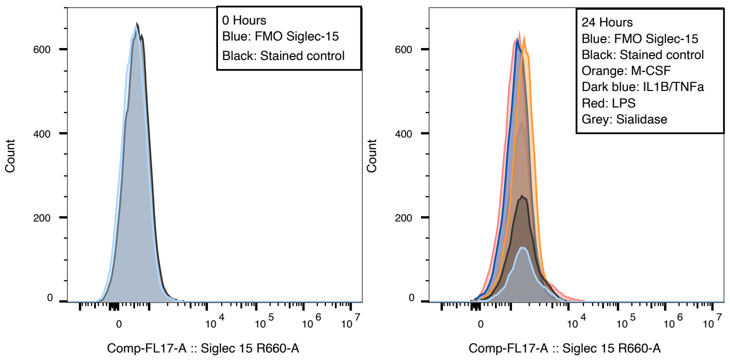


**Figure S3:** We investigated whether Siglec-15 was expressed at 0 and 24hrs but found no evidence of any membrane-associated Siglec-15 expression at these timepoints. The stained samples were compared to the Siglec-15 fluoresence minus one control (FMO).

**Figure S4: Correlation analysis after OA SF stimulation.**

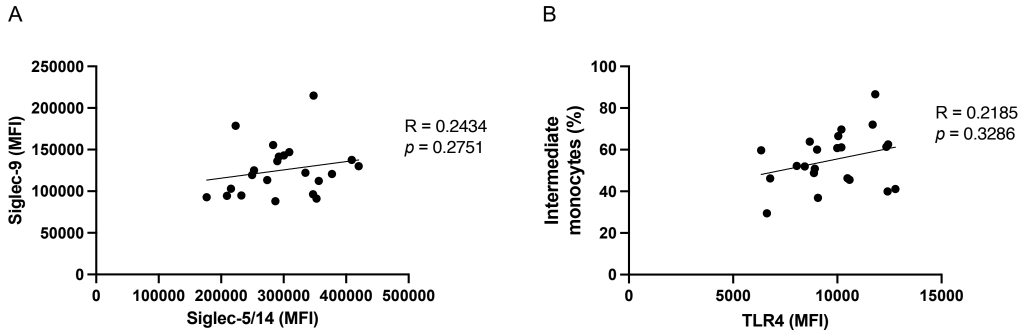


**Figure S4:** Correlation analysis was performed between Siglec-9 and Siglec-5/14 (A) and intermediate monocytes and TLR4 (B) after 24hrs of stimulation with OA SFs. Each data point represents the mean of triplicate samples from one biological replicate. Three biological replicates were tested. Statistical analysis was performed through Spearman's correlation analysis (*$p$<0.05; **$p$<0.01; ***$p$<0.001; ****$p$<0.0001)

**Figure S5: Expression of intermediate monocytes, TLR4, Siglec-5/14, and Siglec-9 at 4hrs of stimulation.**

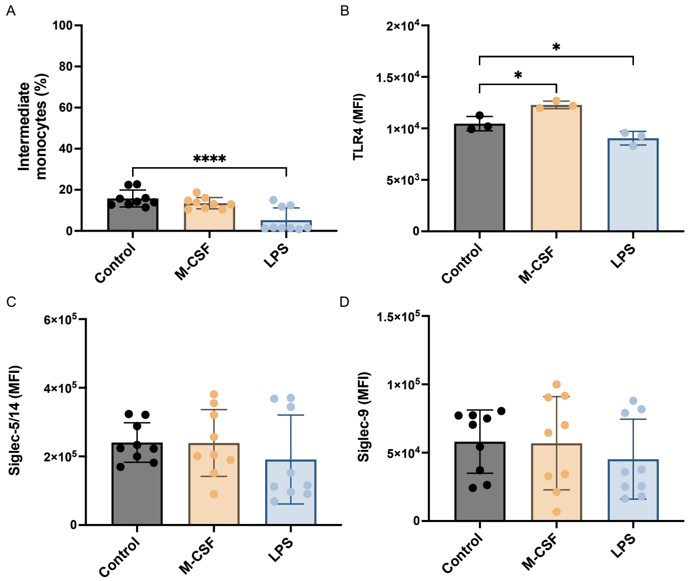


**Figure S5:** The blood monocytes were stimulated with M-CSF and LPS for 4hrs. The percentage intermediate monocytes, based on $CD14^{+}CD16^{+}$ (A), as well as MFI of TLR4 (B), Siglec-5/14 (C), and Siglec-9 (D). Data shown are representative of three biological replicates performed in triplicate, except TLR4, which was performed in one biological replicate. Statistical analysis was performed through one-way ANOVA using Dunnett's multiple comparisons test (*$p$<0.05; **$p$<0.01; ***$p$<0.001; ****$p$<0.0001).

**Figure S6: Expression of TLR4 and Siglec-5 at 24hrs of stimulation with IL1β and TNFα.**

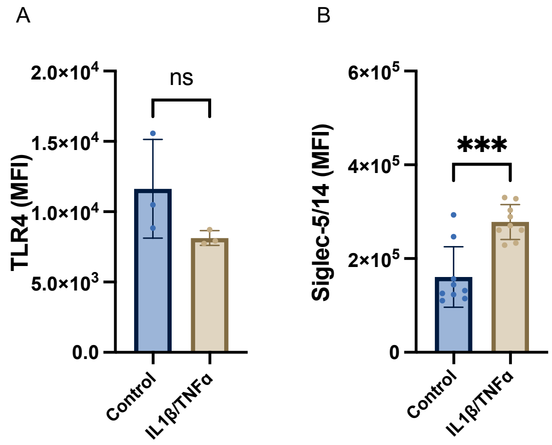


**Figure S6:** The MFI of TLR4 (A) and Siglec-5/14 (B) on monocytes was quantified after 24hrs of stimulation. Data is represented as mean±SD from three technical replicates from one biological replicate for TLR4 and three for Siglec-5. Statistical analysis was performed through one-way ANOVA using Dunnett's multiple comparisons test (**p*<0.05; ***p*<0.01; ****p*<0.001; *****p*<0.0001).

**Figure S7: In-silico protein-protein prediction in AlphaFold3 for direct contact between TLR4 and Siglec-5.**

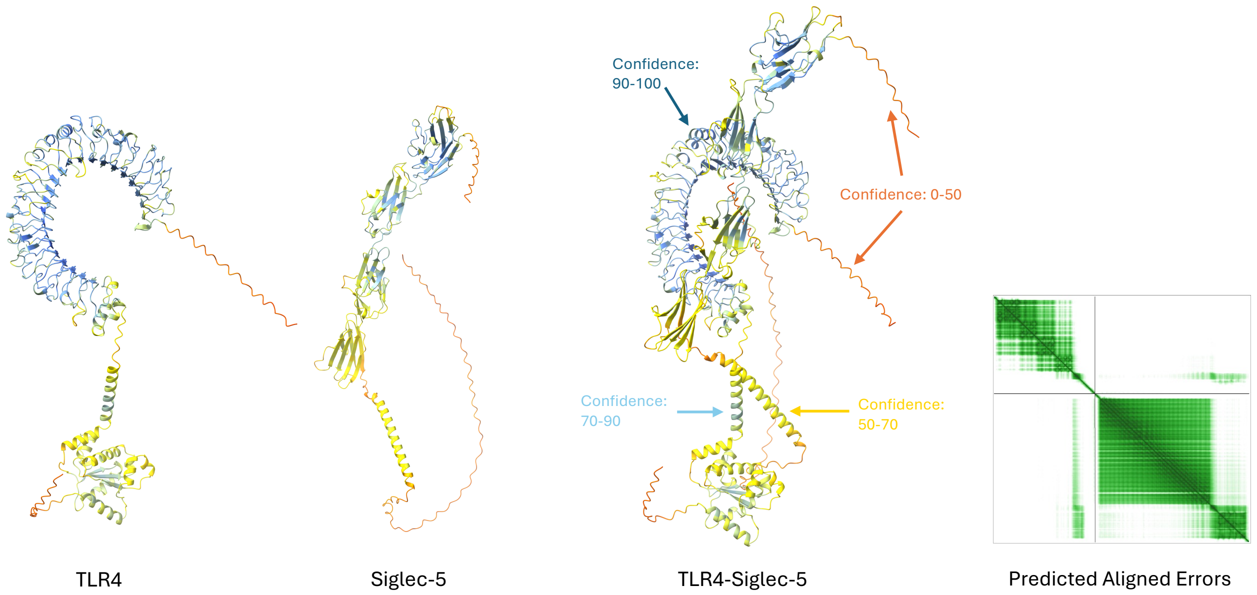


**Figure S7:** The predicted interaction between of TLR4 (O00206) and Siglec-5 (O15389) colored by pLDDT (orange, 0-50; yellow, 50-70; cyan, 70-90; blue, 90-100), were blue indicates high confidence in the prediction and orange color indicates low confidence. The Predicted Alignment Errors (PAE) shows high confidence as green and low confidence as white.

#### Figure S8: TLR4 inhibition after 4hrs.

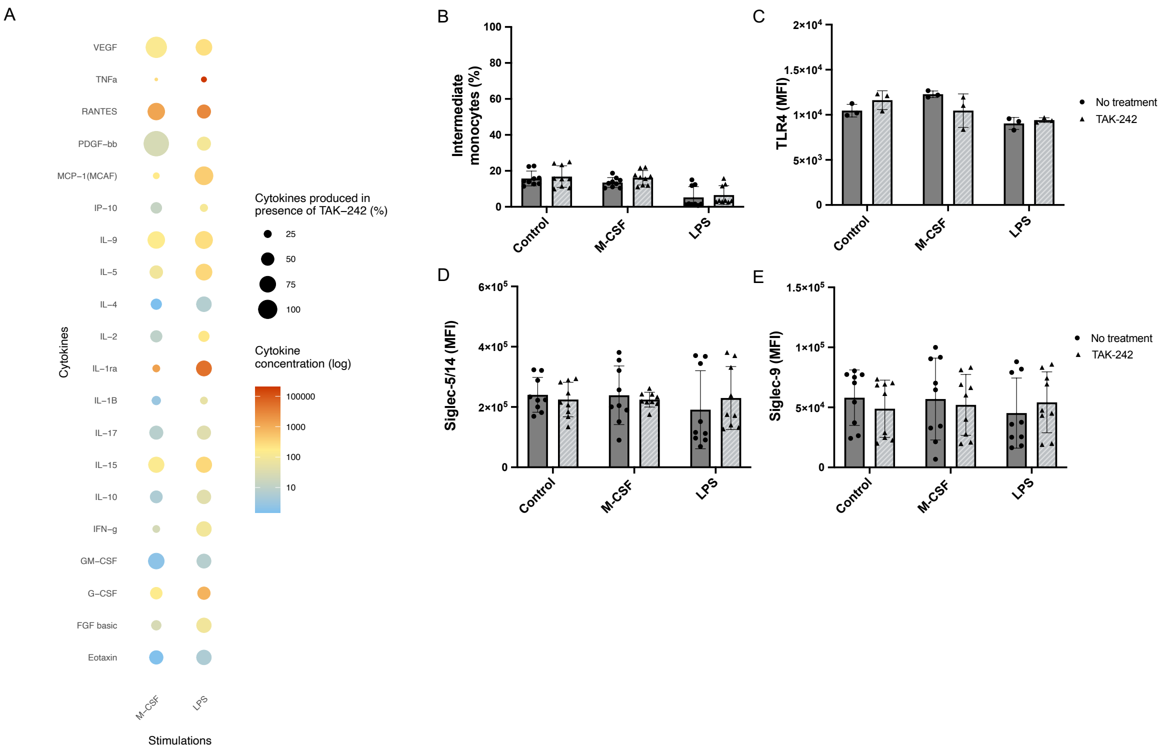


**Figure S8:** Monocytes were subjected to the TLR4 inhibitor, TAK-242, in combination with M-CSF, IL1β/TNFα, LPS, and sialidase for 4hrs. (A) The supernatants of the stimulated monocytes were collected, and panel of cytokines were screened. The effect by TAK-242 on surface receptors was compared to the levels induced by the stimulation without TAK-242. Data is represented as the mean percentage or MFI ± SD (n=3) from three biological replicates. Statistical analysis was performed using two-way ANOVA with Sidak's multiple comparisons test (**p*<0.05; ***p*<0.01; ****p*<0.001; *****p*<0.0001).

**Figure S9: TLR4 inhibition does not limit cytokine production in monocytes stimulated with IL1β/TNFα.**

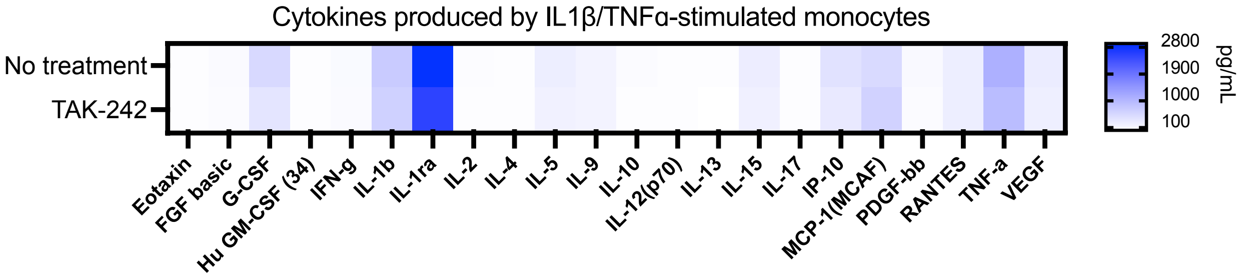


**Figure S9:** A panel of cytokines were screened using the Bio-Plex Pro Human Cytokine Grp I Panel 27-Plex, after 24hrs of stimulating blood monocytes with IL1β and TNFα in presence and absence of TAK-242, this revealed no strong inhibitory effect by TLR4 inhibition. Three technical replicates where pooled from each biological replicate, three biological replicates were used.

**Figure S10: TLR4 inhibition does not affect membrane-expression of Siglec-14.**

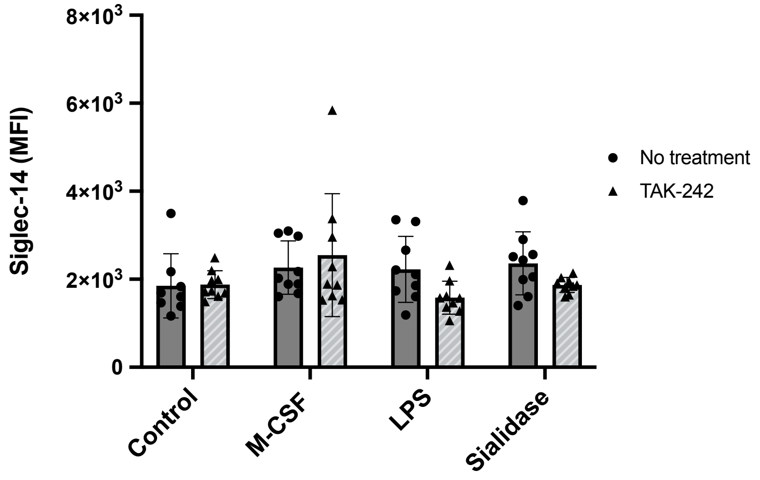


**Figure S10:** Following 24hrs of incubation in presence or absence of the TLR4 inhibitor TAK-242, no significant difference in membrane-expression of Siglec-14 was observed.

**Figure S11: Secondary antibodies do not display unspecific binding.**

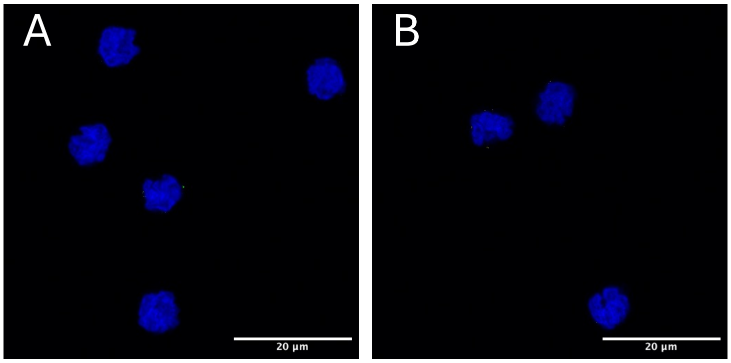


**Figure S11:** (A) Staining with only anti-human Siglec-5 rabbit polyclonal antibody + secondary antibody goat anti-mouse displayed no unspecific binding. (B) anti-human TLR4 mouse monoclonal antibody + secondary antibody goat anti-rabbit displayed no unspecific binding.
